# Vcam1 in endothelial and stromal cells regulates hematopoietic stem cell contact with the niche

**DOI:** 10.1101/2025.08.30.673252

**Authors:** Octavia Santis Larrain, Alice Alhaj Kadour, Sobhika Agarwala, Wantong Li, Bradley W. Blaser, Michael R. Lasarev, Roxana Alexandridis, Anthony Veltri, Khaliun Enkhbayar, Elliott J. Hagedorn, Owen J. Tamplin

## Abstract

Hematopoietic stem and progenitor cells (HSPCs) are essential for differentiation into all blood cell types. In mammals, the interaction between HSPCs and the fetal liver niche during development is critical for stem cell maturation. Integrin alpha 4 (Itga4) on HSPCs and vascular cell adhesion molecule (Vcam1) on niche cells are critical for HSPC colonization of the fetal liver (FL). Itga4 and Vcam1 also function in the zebrafish equivalent of the FL, the caudal hematopoietic tissue (CHT), however, the specific niche cells that express Vcam1 remain unclear. Using multiple approaches, including fluorescent *in situ* hybridization, we found Vcam1 is expressed in endothelial cells (ECs) and mesenchymal stromal cells (MSCs), but not macrophages. Time-lapse live imaging of *itga4* mutants showed the Itga4-Vcam1 axis is required for HSPC retention in the CHT niche, but not homing or lodgment. Our results show that Itga4 on HSPCs and Vcam1 on ECs and MSCs are involved in retention in the CHT niche.

**Summary:** Blood stem cell interaction with the niche microenvironment during development is critical for establishing a robust stem cell pool into adulthood. This study determines the niche cell types that present Vcam1 in the embryo and allow interaction with blood stem cells.

## Introduction

Hematopoietic stem and progenitor cells (HSPCs) are at the top of a hierarchy that produces all blood lineages. Contact with niche cells in the microenvironment is critical for HSPC development in the embryo and homeostasis in the adult (Pinho and Frenette, 2019). In all studied vertebrate embryos, HSPCs emerge from the ventral wall of the dorsal aorta (DA) by budding from the hemogenic endothelium (HE) (Orkin and Zon, 2008). In mammals, HSPCs colonize the fetal liver (FL) and undergo maturation before migrating to the bone marrow (BM) where they persist throughout life (Mikkola and Orkin, 2006). In zebrafish, the equivalent of the FL is the caudal hematopoietic tissue (CHT) where HSPCs reside before migrating to a perivascular niche in the anterior region of the kidney (Agarwala et al., 2022; Murayama et al., 2006; Tamplin et al., 2015). Many different cell types in these embryonic niches, including ECs, MSCs, and immune cells, facilitate colonization of the niche by HSPCs, although the precise mechanisms are not well understood (Agrawal et al., 2024).

Integrins and their ligands are critical for HSPC interaction with the niche during development and into adulthood (Imai et al., 2010; Krenn et al., 2022). One important integrin heterodimer expressed on HSPCs is Itga4/b1 that binds its receptor Vcam1 on niche cells. The Itga4/b1-Vcam1 interaction is highly conserved and critical for colonization of the FL niche in mammals and the CHT niche in zebrafish (Arroyo et al., 1996; Arroyo et al., 1999; Gribi et al., 2006; Hirsch et al., 1996; Koenig et al., 2002; Li et al., 2018; Potocnik et al., 2000; Priestley et al., 2006; Qian et al., 2007; Williams et al., 1991). A recent study of fetal bone marrow showed Vcam1 is expressed in a subset of sinusoidal ECs and is required for HSPC colonization (Weinhaus et al., 2025). In adult BM, Vcam1 is expressed by Lepr+ MSCs, sinusoidal ECs, and osteolineages (Baryawno et al., 2019), and regulates HSPC trafficking into and out of the niche (Craddock et al., 1997; Frenette et al., 1998; Papayannopoulou et al., 1995; Papayannopoulou et al., 1998).

Interestingly, Vcam1 is not only expressed by niche cells but also hematopoietic populations (Ulyanova et al., 2005). Hematopoietic stem cells and leukemic cells express Vcam1 to provide innate immune tolerance in the BM (Pinho et al., 2022). Vcam1+ macrophages interact with Itga4+ primitive erythroid cells in FL erythroblastic islands (Isern et al., 2008; Sadahira et al., 1995). Recently, single cell RNA-sequencing (scRNA-seq) defined a distinct subset of FL macrophages that express Vcam1 (Kayvanjoo et al., 2024). A Vcam1+ embryonic macrophage population is involved in establishment of the adult BM niche and supports MSCs (Percin et al., 2025). In mouse FL, HSPCs develop near macrophages that regulate their differentiation (Gao et al., 2022; Monticelli et al., 2024; Tang et al., 2025). The precise role of Vcam1 and macrophages in the FL is still emerging.

The Itga4-Vcam1 axis is present in mammals and zebrafish, except that zebrafish have *vcam1a* and *vcam1b*, owing to the whole genome duplication in teleosts (Howe et al., 2013). A forward genetic screen discovered *itga4* was required for HSPC colonization of the CHT (Li et al., 2018), confirming Itga4 has a conserved function in HSPC seeding of the mammalian FL (Arroyo et al., 1999). It was also suggested in zebrafish that Vcam1+ macrophages facilitate HSPC lodgment and retention in the CHT (Li et al., 2018). Other work in zebrafish showed macrophages perform quality control on HSPCs in the CHT to prevent inferior clones from contributing to long-term hematopoiesis (Wattrus et al., 2022). We sought to better understand how cell-cell interactions between HSPCs and the niche promotes stem cell maturation in the zebrafish embryo. Our previous correlative light and electron microscopy of the zebrafish CHT and larval kidney identified a typical niche unit made up of an HSPC, multiple ECs, and one MSC (Agarwala et al., 2022; Tamplin et al., 2015). In mouse FL a similar HSPC ‘pocket-like’ unit of ECs, stromal cells, hepatoblasts, and macrophages was described (Gao et al., 2022). Therefore, we hypothesized that a highly evolutionarily conserved Itga4-Vcam1 axis may promote HSPC lodgment in these embryonic niche units. There have been conflicting reports about the cells that express Vcam1 in the zebrafish embryo (Li et al., 2018; Wattrus et al., 2022), so we have used multiple complementary approaches and reanalysis of published dataset to address our hypothesis.

## Results

### Itga4 is required for retention but not homing or EC pocket formation in the CHT

We wanted to further characterize the role of Itga4 in HSPC lodgment in the CHT. A previous study used the *Tg(kdrl:Dendra2)* line to examine HSPC colonization of the CHT in wild-type (WT) and *itga4^cas005^* mutant backgrounds (Li et al., 2018). These experiments depended on green-to-red fluorescent protein photoconversion of HE in the DA, followed by tracking of red DA-derived HSPCs in the unconverted green vasculature of the CHT. To perform similar experiments without the need for photoconversion, we used the alternative approach of combined HSPC-specific (*Tg(Runx:GFP)* (Tamplin et al., 2015)) and EC-specific (*Tg(kdrl:mCherry)* (Chi et al., 2008)) transgenic reporter lines in WT and *itga4^cas005^* mutant backgrounds. We performed time-lapse imaging cycling between multiple *Runx:GFP;kdrl:mCherry* embryos that were WT, heterozygous *itga4^cas005/+^* (Het), or homozygous *itga4^cas005/cas005^.* Time-lapse began from 54 hours post fertilization (hpf) when the CHT is being colonized by HSPCs and continued for >10 hours. As we showed before (Tamplin et al., 2015), Runx:GFP+ HSPCs that lodge in the CHT niche trigger remodeling of ECs to form a surrounding pocket (Fig. 1A; Movie 1; WT or Het *itga4^cas005/+^*). Interestingly, HSPCs in homozygous *itga4^cas005/cas005^* mutants could briefly contact the niche and trigger remodeling, but then shortly after dropped out of the surrounding pocket (Fig. 1A; Movie 2). In this example, the EC pocket is not maintained after egress of the HSPC, suggesting it may be dependent on HSPC contact. In contrast to previous reports that HSPCs could not enter the venous capillaries in *itga4^cas005^* mutants and be surrounded by ECs (Li et al., 2018), we found that Runx:GFP+ HSPCs in *itga4^cas005^* mutants did lodge and were wrapped in an EC pocket in the CHT (Fig. 1A), suggesting that the Itga4-Vcam1 axis is not required to guide HSPCs into the CHT, and instead is required for retention once HSPCs have lodged.

**Figure 1.**
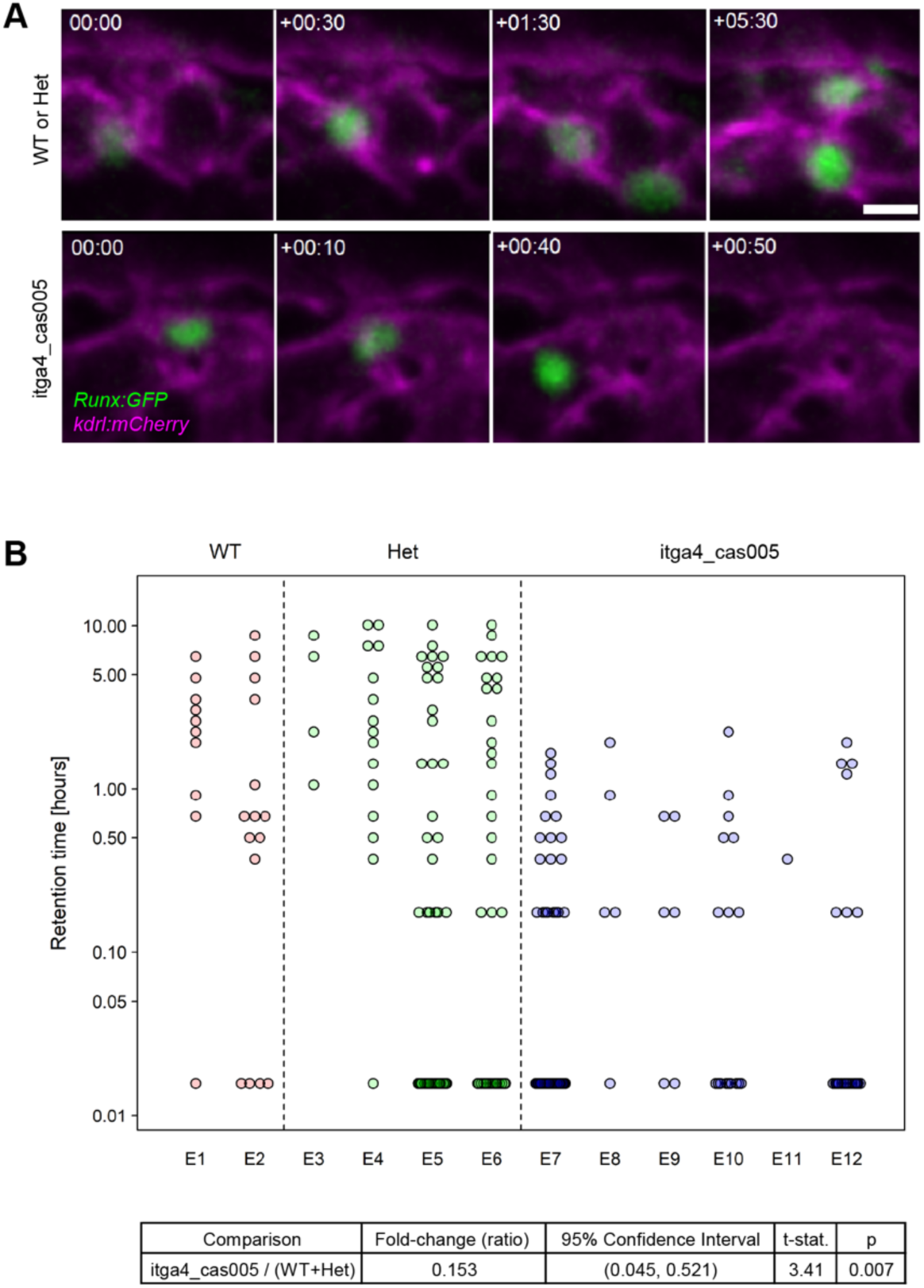
*itga4* is required for retention in the CHT niche but not homing or EC pocket formation. **(A)** Single frames from time-lapse movies showing *Runx:GFP+* HSPCs (green) lodging in *kdrl:mCherry*+ ECs (magenta) of the CHT. Movies start at 54 hpf (time=00:00). Upper row: WT or Het. Lodged HSPC stays in EC pocket >5 hours. Lower row: *itga4^cas005^* mutant. HSPC lodges in CHT and EC pocket forms, but then HSPC drops out of pocket after <40 minutes. Scale bar is 10 microns. Corresponds to Movies 1 and 2. **(B)** Analysis of time-lapse live imaging results. n=12 embryos (2 WT, 4 Het, 6 *itga4^cas005^* mutant); average ∼20 cells recorded/embryo. Retention times ranged from 0-10.83 hours. Not all embryos survived to the end of the experiment. A constant value of 0.015 hours (55 seconds) was added to all retention times to support log-transformation. This value of 55 seconds was used as it was the minimum time noted in similar previously published experiments (Li et al., 2018), allowing comparison between datasets. Increasing all times by 55 seconds leads to retention times ranging from 0.015–10.85 hours. Distribution of retention times in any given embryo are lower in *itga4^cas005^* mutants compared to WT/Het (i.e., retention time for mutants never >2.2 hours; all WT/Het embryos had cells with retention times above >2.2 hours). Retention time did not differ between WT and Het (t_9_ = 0.575, p = 0.580) and were therefore combined into a single group. An overall ANOVA-type test showed retention time differs among WT, Het, and *itga4^cas005^* mutants (F_2,9_ = 5.93, p = 0.023) and justified a further search among genotypes, with suitable correction for multiple testing from these searches. Geometric mean (GM) retention time for *itga4^cas005^* mutants is ∼85% (95% CI: 48–95%) lower than for WT/Het (0.52 vs 0.08 hours; t_10_ = 3.41, p = 0.007). Three experiments were performed and results shown are from one representative experiment.

We performed manual tracking of lodged HSPCs in time-lapse movies to measure the retention time in the CHT (Fig. 1B). WT or Het embryos had sustained lodgment in the CHT, whereas *itga4^cas005^* mutant HSPCs only had brief retention. As a rigorous statistical approach, we used linear mixed-effect models (Pinheiro and Bates, 2009) to test whether retention time differed among genotypes (*itga4^cas005^* vs WT/Het). These models were used to account for the design of each experiment in which the genotype applied to individual embryos, but subsequent recordings of retention time were measured on a collection of cells derived from a given embryo. Genotype was a fixed-effect in all models, while the embryo was considered as a random factor; models further allowed cell-to-cell variation of retention times to differ by genotype. Using this approach we found the geometric mean (GM) of retention time for *itga4^cas005^* mutants (0.08 hours) is approximately 85% (95% confidence interval [CI]: 48-95%; t_10_ = 3.41, p=0.007) lower than the corresponding GM retention time for WT/Het (0.52 hours). We used this same statistical approach to reanalyze previously published raw data of HSPC retention times in time-lapse movies and found the results were very similar to ours (*itga4^cas005^* mutant GM of 0.07 hours vs WT 0.50 hours; Fig. S1A) (Li et al., 2018). Reanalysis of HSPC retention times in *vcam1b^cas011^* mutants were also similar (GM of 0.02 hours; Fig. S1B) (Li et al., 2018). Contrary to published results (Li et al., 2018), reanalysis did not find a significant difference in HSPC retention times in the CHT after nitroreductase-mediated macrophage ablation (Fig. S1B; compare Condition 1 vs 2).

### Polyclonal anti-zebrafish Vcam1b antibody detects embryonic ECs and MSCs by flow cytometry

We wanted to evaluate the utility of a polyclonal anti-Vcam1b antibody to determine cell-type specific expression of Vcam1b protein in the zebrafish embryo. This approach was tested previously (Li et al., 2018), however, that antibody was not available, so we had a polyclonal antibody raised against an epitope of zebrafish Vcam1b (amino acids 18-235); antigen specificity was commercially tested by titration ELISA (Boster Bio cat. no. DZ01199-1). We previously generated a *vcam1b^uwt007^* mutant allele using CRISPR-Cas9 gene editing (Fig. S1C) (Bornhorst et al., 2024). This mutant contains a deletion within the Vcam1b antibody epitope, predicted to create a premature stop codon and truncated protein. We tested the antibody in western blot using brain tissue from a WT adult and detected a single band at the predicted size of zebrafish Vcam1b (88 kDa; A4JYI2_DANRE; Compute pI/Mw (Gasteiger et al., 2005)) (Fig. S1D). The mutant exhibited loss of HSPC marker *cmyb* in the CHT at 5 dpf, consistent with the previously published *vcam1b^cas011^* mutant allele (Li et al., 2018) (Fig. S1E). We used brain tissue from *vcam1b^uwt007^* adult zebrafish to again test the Vcam1b antibody in western blot but did not detect a significant reduction in Vcam1b protein levels (Fig. S1D). Our results indicate the polyclonal anti-Vcam1b antibody detects protein similar in size to endogenous Vcam1b, with the caveat it may also detect other similar proteins in zebrafish like Vcam1a.

Using whole mount *in situ* hybridization (WISH) we confirmed the endogenous expression pattern of *vcam1b* matched published reports of expression in the brain, heart, and CHT (Fig. S1F; zfin.org (Thisse et al., 2008)). Using fixed transgenic embryos, whole mount immunofluorescence (IF) was performed against mCherry in *ECs (kdrl:mCherry)* and GFP in macrophages (*mpeg1:GFP*). However, we could not reliably use our polyclonal anti-Vcam1b antibody due to non-specific staining that was inconsistent with endogenous expression (e.g., ectopic expression in intersomitic vessels and neural tube; Fig. S1G). For that reason, we could not confirm the previously published report of Vcam1b expression in *mpeg1:GFP*+ macrophages in the CHT that used whole mount IF and fixed embryos (Li et al., 2018). It has been notoriously difficult to raise highly specific antibodies against zebrafish proteins, particularly those that work in fixed tissues (Staudt et al., 2015). One possible reason is the glycans and disulfide bonds present on zebrafish proteins that are not present in the peptides synthesized as antigens (Traver et al., 2003).

Although the anti-Vcam1b antibody was not specific in whole mount fixed embryos, we wanted to test it in live embryos by intravascular injection as a dye-conjugated antibody (Fig. S2A), as was previously performed (Li et al., 2018). We used intravascular retro-orbital injection into *mpeg1:GFP;kdrl;mCherry* embryos with control IgG-647 or conjugated anti-Vcam1b-647 antibody. We co-injected blue dextran to broadly label the inner lumen of the vasculature. We found there was widespread vascular endocytosis of IgG-647, anti-Vcam1b-647, and blue dextran in the CHT (Fig. S2B). We tested a range of doses of IgG-647 and found a dose-dependent increase in vascular endocytosis in the CHT (Fig. S2C), including doses used previously (Li et al., 2018). Lastly, we performed time-lapse live imaging of IgG-647 or anti-Vcam1b-647 and blue dextran injected into *mpeg1:GFP;kdrl;mCherry* embryos (Fig. S2D,E; Movies 3,4). We observed migratory *mpeg1:GFP+* macrophages that had taken up cellular debris labeled with IgG-647, anti-Vcam1b-647, blue dextran, and/or mCherry from *kdrl;mCherry*+ ECs. Our results suggest injection of conjugated antibody into the circulation of the live zebrafish embryo generates non-specific labeling that together with cellular debris is taken up by macrophages. This is consistent with the observation that macrophages patrolling the CHT can take up small portions of cellular material or can fully engulf cells (Wattrus et al., 2022). Our results show that any exogenously injected conjugated antibody or dextran, as well as endogenously expressed fluorescent proteins, can be taken up and carried by macrophages in the CHT. This indicates the technique of intravascular injection in the live zebrafish embryo must be carefully controlled to ensure the specificity of labeling.

As an alternative to whole mount IF of fixed embryos and intravascular injection of live embryos, we tested the anti-Vcam1b-647 antibody using flow cytometry. We dissociated 54 hpf embryos and stained single cell suspensions with anti-Vcam1b-647 antibody. We used the following transgenic reporter lines: EC-specific *kdrl:mCherry* (Whitesell et al., 2019); MSC-specific *ET37:GFP* (Murayama et al., 2023; Parinov et al., 2004); *cxcl12:dsRed* (Glass et al., 2011); macrophage-specific *mpeg1:GFP* (Kuil et al., 2020). We quantified cells that were positive for both transgenic reporter fluorescent protein and Vcam1b-647, using IgG-647 as a negative control (Fig. 2A). Unlike intravascular injection of IgG-647 in the embryo that generated non-specific signal from vascular endocytosis (Fig. S2), control IgG-647 in flow cytometry allowed discrimination and gating of cells that were Vcam1b-647 positive or negative (Fig. 2A).

**Figure 2.**
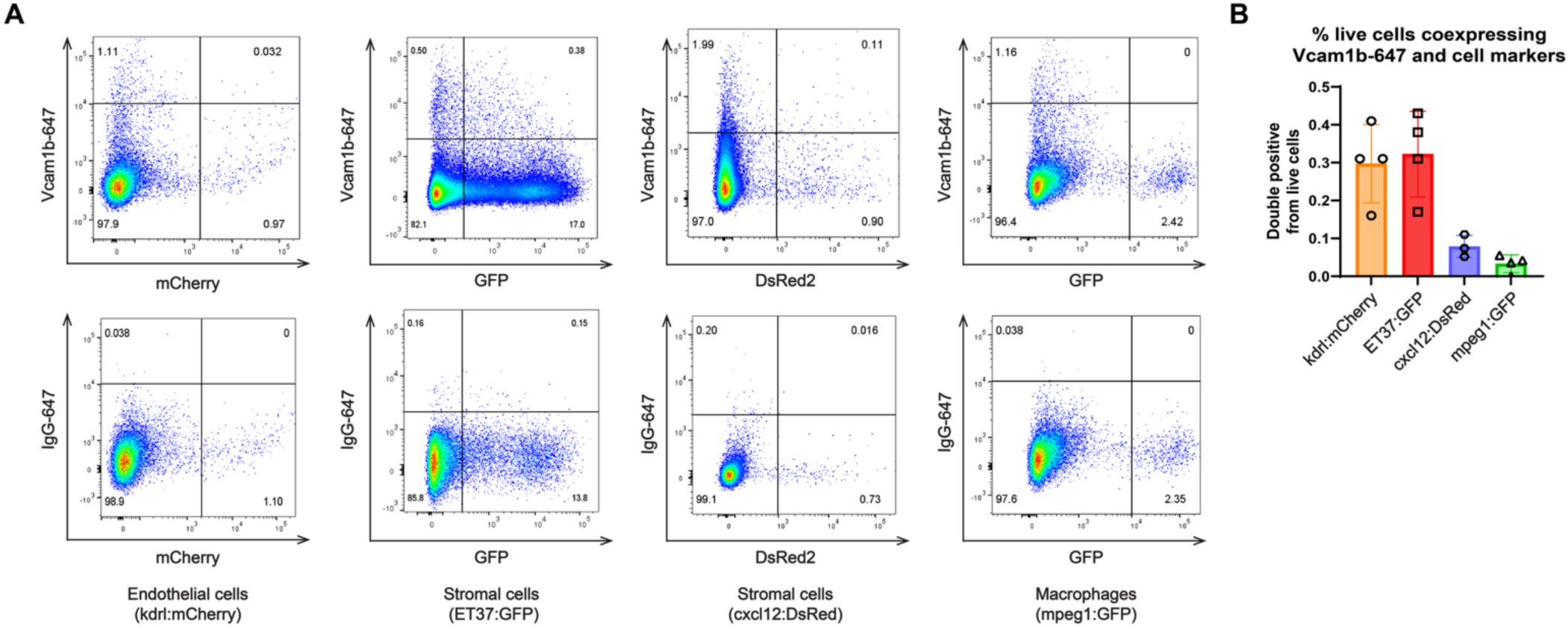
Polyclonal anti-Vcam1b antibody detects ECs and MSCs using flow cytometry. **(A)** Upper: flow cytometry of Vcam1b-647+ cells from pooled dissociated 54 hpf embryos: ECs (*kdrl:mCherry*); MSCs (*ET37:GFP* and *cxcl12:dsRed*); macrophages (*mpeg1:GFP*). Lower: IgG-647 gating control. **(B)** Quantification of double-positive cells in (A) as a percentage of total live cells.

Quantification of double positive cells found a proportion of ECs and MSCs but not macrophages were bound by the anti-Vcam1b antibody (Fig. 2B). If anti-Vcam1b-647 antibody is marking MSCs, then it is not surprising the proportion of double positive *cxcl12:dsRed+* Vcam1-647+ cells is lower than *ET37:GFP+* Vcam1-647+ because the *cxcl12:dsRed* reporter line is less specific and marks cell types other than MSCs (Glass et al., 2011).

### Sorted Vcam1b+ embryonic ECs and MSCs express endogenous *vcam1b* transcript

Given the importance of the Itga4-Vcam1 axis for HSPC lodgment in the CHT, we wanted to define all niche cells that express Vcam1b and thereby identify any potentially important cell-cell interactions involved in developmental maturation of HSPCs. To confirm that cells detected by anti-Vcam1b-647 antibody and flow cytometry also expressed endogenous *vcam1b* transcript, we sorted these cells for scRNA-seq (Fig. 3A,B). UMAP analysis and unsupervised Leiden clustering of the scRNA-seq data revealed 10 distinct clusters (Fig. 3C). We detected *vcam1b* expression in almost all the clusters (Fig. 3D,E), including those that expressed markers for MSCs (*cspg4*, *cxcl12a*, *krt4*, *lepr*, *pdgrfrb*; clusters 3 and 4) and ECs (*kdrl*; cluster 8). Expression of *vcam1b* in neuronal cells (*elavl3*; clusters 1, 2, and 10), was consistent with our own (Fig. S1F) and published WISH data (Thisse and Thisse, 2008a; Yang et al., 2015), and published scRNA-seq data (Fig. S3). Although a small number of macrophages were captured in our cell sorts for scRNA-seq, as indicated by a *mpeg1.1+* population (cluster 9), this cluster was negative for *vcam1b*. The related *vcam1a* gene was also not expressed in *mpeg1.1+* cluster 9 (Fig. 3E). These results suggest a polyclonal conjugated anti-Vcam1b-647 antibody can capture *vcam1b+* ECs, MSCs, and neuronal cells.

**Figure 3.**
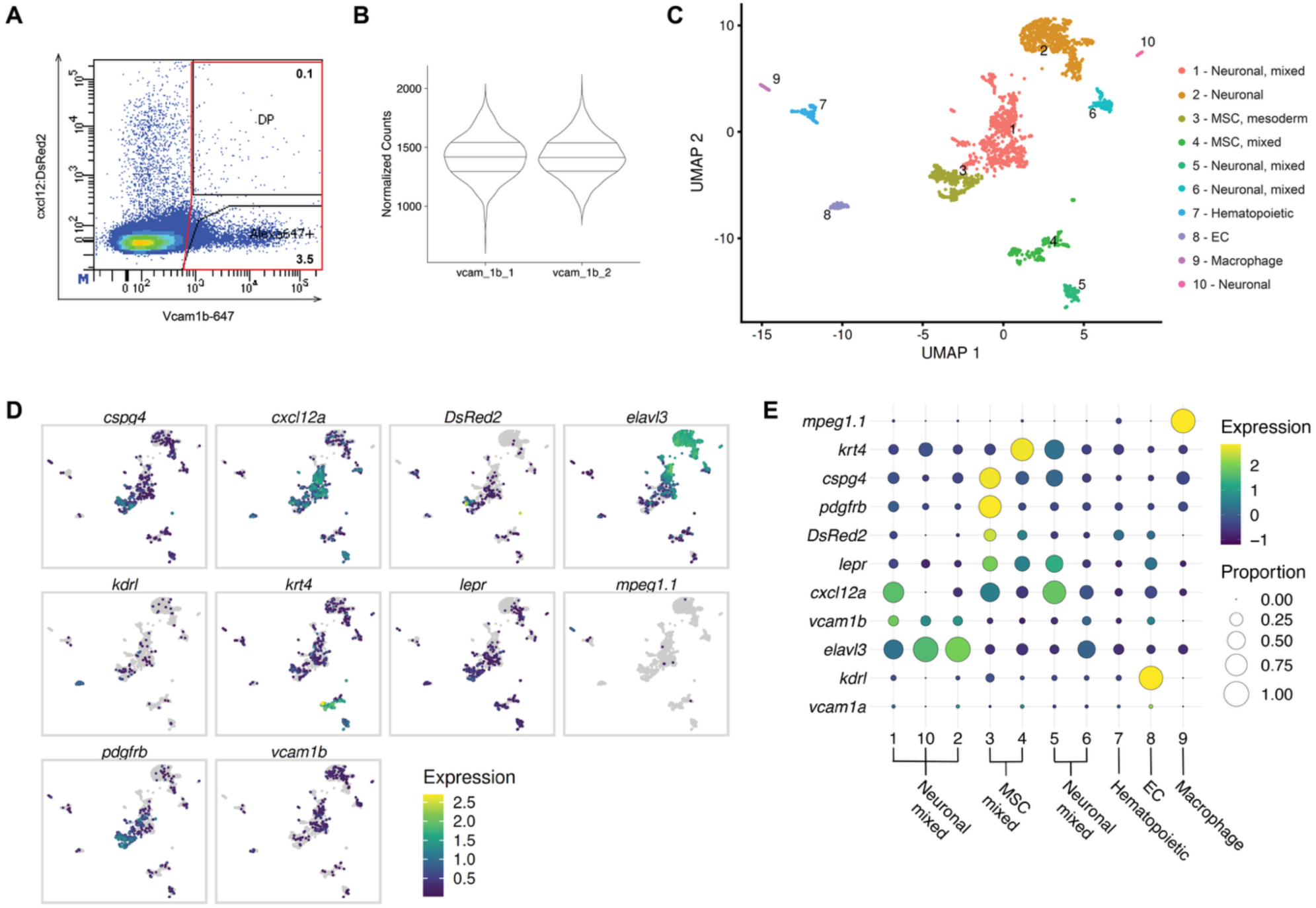
Sorted Vcam1b+ cells express markers of ECs, MSCs, and neuronal cells. **(A)** Flow cytometry gate (red) used to sort all Vcam1-647+ cells from dissociated 54 hpf *cxcl12:DsRed2+* and *casper* embryos. **(B)** Violin plots showing normalized gene counts/cell of 2 technical replicates (vcam1b_1 and vcam1b_2). **(C)** UMAP scRNA-seq Leiden clustering of all Vcam1b-647+ sorted cells. Automated annotation of clusters was followed by manual curation based on known markers. “Mixed” indicates different cell types were annotated to the same cluster and could not be easily resolved. **(D)** Leiden clusters that express *vcam1b* and candidate markers for MSCs (*cspg4, cxcl12a, DsRed2* from *cxcl12:dsRed, krt4, lepr, pdbfrb*), neuronal cells (*elavl3*), ECs (*kdrl*), and macrophages (*mpeg1.1*). **(E)** Bubble plot showing the same clusters and marker genes as in (D), as well as *vcam1a*. Bubble size represents proportion of cells expressing the marker, and color indicates per-cell expression.

### *vcam1b* transcript is expressed in ECs and MSCs but not macrophages

We looked to published scRNA-seq datasets (Farnsworth et al., 2020; Lange et al., 2023; Speir et al., 2021) to further validate our findings that zebrafish ECs and MSCs express *vcam1b*. We found clusters of *cspg4*+ MSCs and *kdrl*+ ECs expressed *vcam1b*, while in *mpeg1.1*+ macrophage clusters *vcam1b* was absent (Fig. S3A,B). Considering that transcripts expressed at low levels may be below the level of detection by scRNA-seq, we reanalyzed published bulk RNA-seq datasets of sorted zebrafish ECs (Lawson et al., 2020; Whitesell et al., 2019), mesenchymal cells (Lawson et al., 2020; Whitesell et al., 2019), and macrophages (Kuil et al., 2020; Rougeot et al., 2019; Theodore et al., 2017) (Fig. 4). Sorted ECs expressed high levels of EC-specific marker *kdrl* and *vcam1b* compared to negative controls, while *cxcl12b* was expressed in both positive and negative populations (Fig. 4A), indicative of its expression in multiple tissues (Knaut et al., 2005). Sorted mesenchymal populations, such as vascular mural and smooth muscle cells, epidermis, cartilage, and pericytes, had high levels of *cxcl12b*, lower levels of *vcam1b*, and varied expression levels of *pdgfrb* dependent on the population that was sorted (Fig. 4B). Sorted macrophages from *mpeg1:GFP*, *mpeg1:mCherry*, and *mpeg1:GAL4;UAS:Kaede* transgenic lines (Ellett et al., 2015; Kuil et al., 2020; Rougeot et al., 2019; Theodore et al., 2017), and at different stages (28, 50, 72, 120, 144 hpf), showed high levels of macrophage-specific markers *mpeg1.1* and *mfap4.1*, intermediate levels of *irf8*, and no expression of *vcam1b* (≤5 FPKM; Fig. 4C).

**Figure 4.**
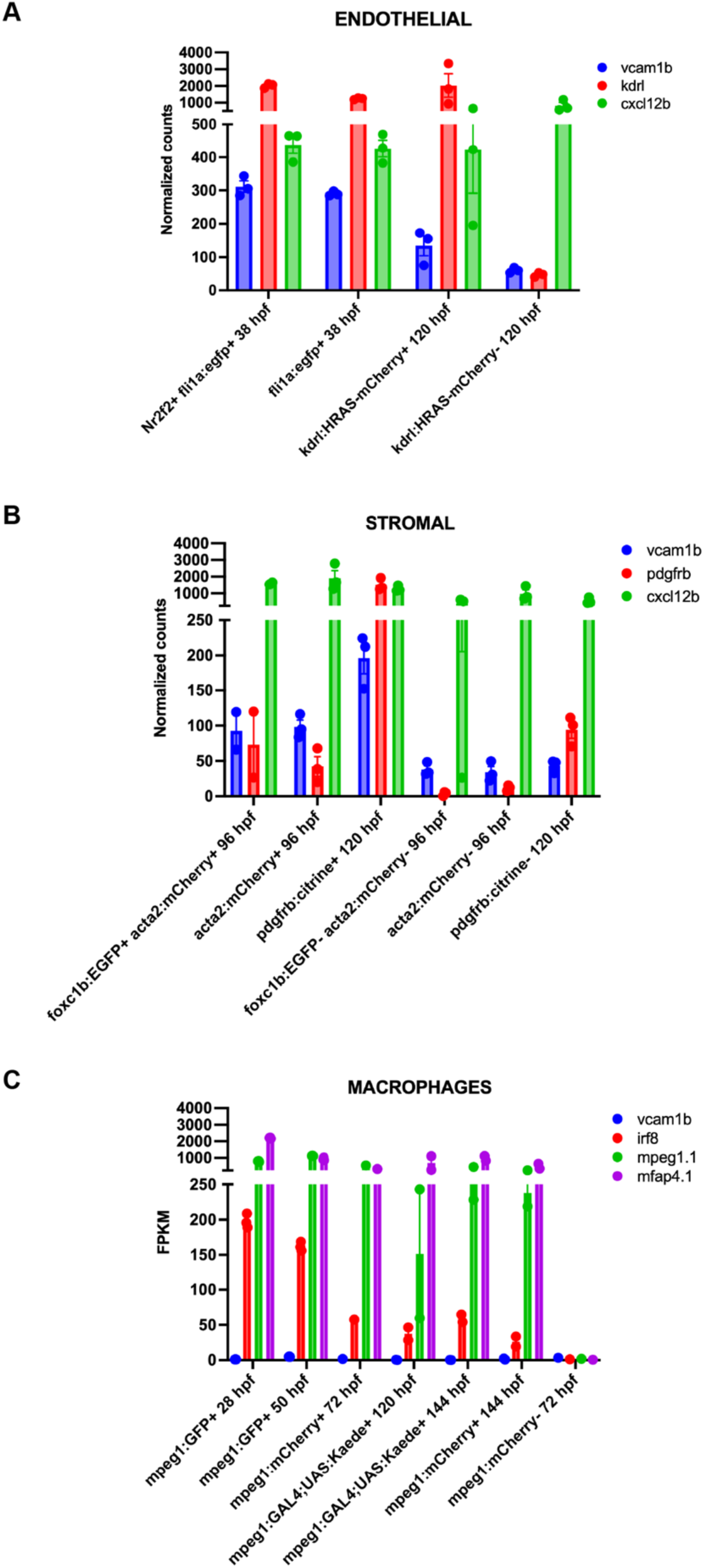
Reanalysis of published bulk RNA-seq datasets shows *vcam1b* expressed in sorted pools of ECs and MSCs but not macrophages. Bulk RNA-seq data shows *vcam1b* is expressed in ECs and MSCs but not macrophages. **(A)** EC-specific marker *kdrl* is enriched in sorted ECs (datasets: *Nr2f2+/fli1a:egfp+* (Lawson et al., 2020); *fli1a:egfp*+ (Lawson et al., 2020); *kdrl:HRAS-mCherry+* and negative (Whitesell et al., 2019)). **(B)** *vcam1b* and MSC markers *pdgfrb* and *cxcl12b* are expressed in sorted MSCs (datasets: *foxc1b:EGFP+;acta2:mCherry+* and negative (Whitesell et al., 2019); *acta2:EGFP+* and negative (Whitesell et al., 2019); *pdgfrb:citrine+* and negative (Lawson et al., 2020)). **C)** Macrophage markers *mpeg1.1*, *mfap4.1*, and *irf8* were expressed in sorted macrophages but *vcam1b* was not (≤5 FPKM; datasets: *mpeg1:GFP+* (Kuil et al., 2020), *mpeg1:mCherry+* and negative (Theodore et al., 2017), *mpeg1:mCherry+* (Rougeot et al., 2019); *mpeg1:GAL4;UAS:Kaede+* and *mpeg1:mCherry+* (Rougeot et al., 2019)). Developmental stages of each population shown on graph.

Our own and published WISH data showed *vcam1b* is expressed in the CHT, however, these experiments did not identify the cell type (Fig. S1F; zfin.org (Thisse et al., 2008)). Previous reports of Vcam1b expression were solely dependent on detection by antibody in fixed whole mount IF or intravascular injection (Li et al., 2018), but did not consider high resolution expression data for endogenous *vcam1b* transcript. To determine the spatial distribution of endogenous *vcam1b* transcript in the zebrafish embryo at 54 hpf, we used a custom whole mount fluorescent *in situ* hybridization probe (FISH; RNAscope (Wang et al., 2012)), together with whole mount IF of transgenic reporter lines to mark specific populations: EC-specific *kdrl:mCherry+*; MSC-specific *ET37:GFP+*; macrophage-specific *mpeg1:GFP+* (Fig. 5A). RNAscope allows subcellular resolution of single transcripts, and together with fluorescent protein expressed by transgenic reporter lines, we scored transcripts within the boundaries of the above cell types. Image analysis software (Imaris) allowed us to create “surfaces” to score *vcam1b* transcript inside or outside of a cell boundary. Converting *vcam1b* transcripts to “spots” we quantified the number of spots inside surfaces and consistently observed *vcam1b* transcripts within *kdrl:mCherry+* ECs and *ET37:GFP+* MSCs, but not *mpeg1:GFP+* macrophages (Fig. 5B). Together, our data shows that the Itga4-Vcam1 axis in the CHT likely functions between Itga4+ HSPCs and Vcam1b+ MSCs and ECs.

**Figure 5.**
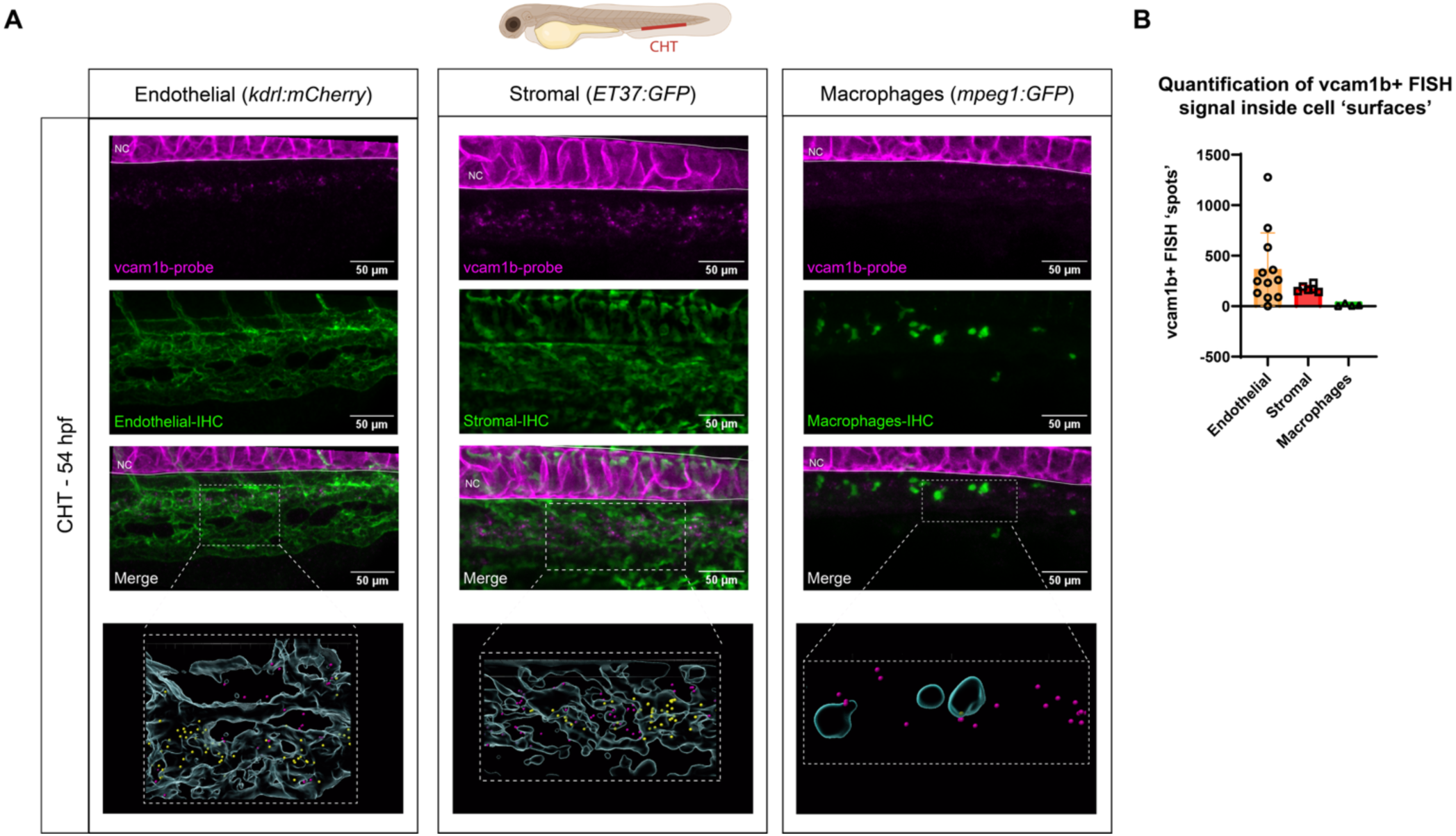
Combined FISH and IF shows *vcam1b* transcripts inside ECs and MSCs but not macrophages. **(A)** At top: diagram of embryo with CHT region marked in red. Representative maximum intensity projections of confocal z-stacks (20x magnification) from the CHT of fixed 54 hpf embryos. FISH showing *vcam1b* transcripts (magenta) with IF (green) to label fluorescent proteins expressed by transgenic reporters: endothelial (*kdrl:mCherry*); stromal (*ET37:GFP*); macrophages (*mpeg1:GFP*). The bottom panel shows 3D volume rendering (Imaris) of transgenic reporters and *vcam1b* transcripts shown as “spots” that are outside (magenta) or inside (yellow) cell “surfaces” (cyan). Note: non-specific signal trapped in notochord at top of images (magenta). **(B)** Quantification of *vcam1b* FISH signal inside cell “surfaces” shown in (A). endothelial (*kdrl:mCherry*; n=12 embryos); stromal (*ET37:GFP*; n=6 embryos); macrophages (*mpeg1:GFP*; n=4 embryos).

## Discussion

A functional genetic role for *itga4* and *vcam1b* in the lodgment of HSPCs in the CHT has been established (Li et al., 2018). However, the specific cell types expressing Vcam1b within the CHT remains unclear. Several cell types, including ECs, MSCs, and macrophages, are potential candidates based on Vcam1 expression in mammalian FL (Jaspers et al., 1995; Kayvanjoo et al., 2024; Koenig et al., 2002; Percin et al., 2025; Schweitzer et al., 1996; Sugiyama et al., 2010; Tada et al., 2006; Ulyanova et al., 2005). We sought to better define the Itga4-Vcam1 axis in the zebrafish CHT. We raised a polyclonal anti-Vcam1b antibody that was tested with indirect ELISA, western blot, fixed embryo whole mount IF, intravascular embryo injection, and flow cytometry. The ELISA confirmed the antibody’s affinity for the antigen was strong, and western blot detected a single band at the predicted size for zebrafish Vcam1b (88 kDa; Fig. S1D). Yet western blot of *vcam1b* mutant tissue still showed a band at the same size (Fig. S1D), suggesting the antibody may bind to another similar protein. We did not have the anti-Vcam1b antibody available from previous studies (Li et al., 2018), so could not reproduce results from whole mount IF and intravascular embryo injection that showed *mpeg1:GFP*+ macrophages were Vcam1b positive and MSCs were Vcam1b negative. Our anti-Vcam1b antibody was non-specific in whole mount IF and intravascular injections (Fig. S1G, S2), but we were able to use it in FACS to isolate ECs, MSCs, and neuronal cells that by scRNA-seq expressed endogenous *vcam1b* transcript. This was consistent with publicly available scRNA-seq (Fig. S3) and bulk RNA-seq (Fig. 4) datasets that showed ECs and MSCs, but not macrophages, expressed *vcam1b* transcript. Finally, we showed using FISH (RNAscope) with a custom *vcam1b* probe that transcript was present within the cell boundaries of *kdrl:mCherry*+ ECs, *ET37:GFP*+ MSCs, but not *mpeg1:GFP*+ macrophages. Our results suggest that in zebrafish *vcam1b* functions in ECs and MSCs in the CHT niche, but is not expressed in embryonic macrophages, unlike mammals that have a well-characterized VCAM1+ macrophage population (Gao et al., 2022; Kayvanjoo et al., 2024; Monticelli et al., 2024; Percin et al., 2025; Tang et al., 2025).

Our results leave questions open about the function of the Itga4-Vcam1 axis in HSPC homing, lodgment, and retention in the embryonic CHT niche. Our live imaging of Runx:GFP+ HSPCs interacting with the CHT vasculature suggests that in the absence of *itga4* guidance, lodgment, and EC pocket formation proceeds normally, but retention in the niche fails. Based on these observations, together with our findings that zebrafish embryonic macrophages do not express *vcam1b*, the question remains for the role of macrophages in the CHT. We see macrophages actively patrolling the dorsal region of the CHT and spending significant time interacting with HSPCs (Fig. S2, Movies 3 and 4), as was observed by others (Li et al., 2018; Wattrus et al., 2022). We also observed these macrophages have high phagocytic activity (Fig. S2, Movies 3 and 4), like mammalian embryonic macrophages (Percin et al., 2025). This is consistent with the function of CHT macrophages performing quality control on HSPCs by either taking up small portions of cells or engulfing them completely (Wattrus et al., 2022). Macrophage depletion experiments have raised additional questions. Macrophage depletion does not affect HSPC colonization and retention of the FL niche, but it does impact HSPC lineage commitment and differentiation (Kayvanjoo et al., 2024; Monticelli et al., 2024). Macrophages in the adult BM niche facilitate HSPC retention (Chow et al., 2011; Winkler et al., 2010), as do VCAM1+ macrophages in the spleen (Dutta et al., 2015). Using an alternative statistical method to reanalyze published data (Li et al., 2018), we found nitroreductase-mediated ablation of macrophages did not change HSPC retention time in the CHT (Fig. S1B). Macrophage depletion did reduce overall definitive hematopoiesis in the CHT (Li et al., 2018; Travnickova et al., 2015), but that was the result of timed clodronate or nitroreductase ablation experiments that may leave contaminating macrophage debris in the microenvironment that has a negative effect on hematopoiesis. Alternative genetic approaches to remove macrophages, either by *irf8* or *cebpa* morpholino knockdown, had either minimal or no reduction in CHT hematopoiesis macrophages (Li et al., 2014; Wattrus et al., 2022; Yuan et al., 2019), further supporting the model that HSPCs can lodge in the CHT in the absence of macrophages. Further investigation will be needed to determine if the CHT “hot-spots” that were observed for HSPC interaction with the niche (Li et al., 2018), may have another explanation, such as MSCs that we showed anchor HSPCs (Agarwala et al., 2022; Tamplin et al., 2015), or subsets of ECs that form specialized niches (Hagedorn et al., 2023). The latter would be consistent with the specialized subsets of ECs that are observed in adult mouse BM (Wu et al., 2024; Zhang et al., 2021), and VCAM1+ sinusoidal ECs in the fetal BM niche (Weinhaus et al., 2025).

## Materials and methods

### Zebrafish lines and husbandry

Zebrafish were handled at the University of Wisconsin-Madison in accordance with Institutional Animal Care and Use Committee Guidelines. Zebrafish were mated, staged, and raised as previously described (Westerfield, 2000). Embryos were maintained at 28.5°C in E3 medium (5 mM NaCl, 0.17 mM KCl, 0.33 mM CaCl_2_, 0.33 mM MgSO_4_) with methylene blue. The *vcam1b^uwt007^* mutant allele was generated previously (Bornhorst et al., 2024) by CRISPR-Cas9-mediated gene editing. Briefly, a sgRNA targeting the *vcam1b* gene (sequence: GCGCGGCAGCTCCAGCGTGT; Synthego) was co-injected with EnGen Spy Cas9 NLS protein as an RNP complex into one-cell stage zebrafish embryos, as previously described (Burger et al., 2016). Adult fish derived from these embryos were screened for mutations by next-generation sequencing (NGS) of PCR products using Illumina MiSeq (Forward primer: ACACTCTTTCCCTACACGACGCTCTTCCGATCTCGGTCCAGACAAACTTCACC; Reverse primer: GTGACTGGAGTTCAGACGTGTGCTCTTCCGATCTTGCAACTGAACAAGCATGAA), and CRISPResso analysis (Clement et al., 2019; Pinello et al., 2016). A founder with a 7 bp deletion within the *vcam1b* coding sequence was identified. Subsequent genotyping of offspring was performed by PCR amplification of the *vcam1b* locus (same primers as above), followed by restriction digest with the Dra-I enzyme to differentiate between homozygous, heterozygous, and wild-type individuals. The *itga4^cas005^* mutant allele was discovered in a forward genetic screen and genotyped as in previously published methods (Li et al., 2018). Transgenic lines used were as follows: *Runx:GFP* (Tamplin et al., 2015); *kdrl:mCherry* (aka *s896Tg* or *Tg(kdrl:Hsa.HRAS-mCherry)*) (Chi et al., 2008); *mpeg1:GFP* (aka *Tg(mpeg:gfp)* or *Tg(mpeg1:EGFP)gl22)*) (Ellett et al., 2011); *ET37:GFP* (aka *Et(krt4:EGFP)*) (Parinov et al., 2004); *cxcl12:dsRed* (aka *Tg(cxcl12a:DsRed2)* or *Tg(sdf-1a:DsRed2)* (Glass et al., 2011). Lines were maintained on AB wild-type (WT) or *casper* (White et al., 2008) backgrounds.

### Time-lapse live imaging of CHT colonization

This was performed as previously described (Tamplin et al., 2015). Briefly, transgenic zebrafish lines were crossed, staged, and selected by fluorescence microscopy. Zebrafish embryos were mounted in multi-well MatTek glass bottom dishes (No. 1.5 cover slip) with 1% LMP agarose, covered with E3 media, tricaine to immobilize, and 0.003% PTU (1-Phenyl-2-thiourea) to block melanogenesis. Time-lapse was started at 54 hpf in an incubated chamber at 28.5°C and performed up to ∼10 hours in duration. Live confocal microscopy was performed using a Yokogawa spinning disk and Nikon inverted Ti microscope. Objectives lens was a Nikon 20x Plan-Apo DIC N.A. 0.75. Image acquisition was done with dual Andor iXon x3 EM-CCD cameras (512×512 pixels) or dual Hamamatsu Quest cameras. Multiple embryos were imaged within a 3– 10 min interval using a moving XY stage, as well as acquisition of Z stacks through the entire CHT in multiple fluorescent channels. The instrument was controlled using Nikon NIS-Elements software. After time-lapse imaging embryos were recovered for genotyping as previously described for WT, Het, or homozygous *itga4^cas005^* mutants (Li et al., 2018). Image processing and rendering was done using Imaris (Bitplane) and/or NIS-Elements (Nikon). Cell tracking to measure duration of HSPC lodgment in the CHT was done using the ImageJ/Fiji and the MTrackJ plugin (Meijering et al., 2012; Schindelin et al., 2012).

### Statistical analysis of HSPC retention time in time-lapse movies

Retention time (in hours) was log transformed prior to analysis to improve symmetry and stabilize variation in the retention times across treatments. Means calculated on log-transformed data represent geometric means (GM) after back transformation to the original scale. At the same time, differences between means of log-transformed data become fold-changes (ratios) involving these geometric means after back transformation. Tukey’s method was used to adjust p-values for the set of pairwise comparisons in each experiment. Analyses were performed using R (v 4.4.0) (R_Core_Team, 2021) using the nlme package (Pinheiro et al., 2023).

### Antibodies and additional reagents

Polyclonal rabbit anti-Vcam1b antibody was raised to target the epitope between amino acids 18 and 235 (zebrafish protein sequence UniProt A5PMX0_DANRE) using a custom service from Boster Biological Technology (Cat. No. DZ01199-1; antigen specificity tested by titration ELISA). Conjgated anti-Vcam1b-iFluor-647 antibody (Cat. No. DZ01199-1-iFluor647; Boster Bio) and control Goat Anti-Rabbit IgG-Alexa Fluor 647 (Cat. No. ab150079; Abcam) were used for intravascular embryo injections and flow cytometry. Blue dextran (Cascade Blue, 10,000 MW, Anionic, Lysine Fixable; Cat. No. D1976; Invitrogen) was used to label vessel lumen in intravascular embryo injections.

### Protein extraction and western blot

Adult fish brains were homogenized in 2X Cell lysis Buffer (Cell Signaling Technology; Cat. No. 9803S) with PMSF (Cell Signaling Technology; Cat. No. NC0417414). Protein extracts were prepared with loading buffer (1X) and separated by SDS-PAGE on 4-12% Bis-Tris gels (cat no NW04120). One gel was stained with Blue Coomassie to determine protein integrity, and a duplicate gel was transferred to PVDF membranes. Membranes were probed with anti-Vcam1b antibody (1:500; Cat. No. DZ01199-1; Boster Bio) and anti-β-actin (1:1000; Cell Signaling Technology, Cat. No. 4967S) antibodies, followed by HRP-conjugated secondary antibody (Goat anti-Rabbit HRP, cat no 65-6120). Protein bands were detected using chemiluminescence and visualized with an LI-COR imager. Quantification was performed in Fiji/ImageJ.

### Whole mount immunofluorescence (IF)

Embryos were fixed overnight in AB buffer containing 4% PFA in 2X fix buffer (234 mM sucrose, 0.3 mM CaCl2, and 1X PBS). Post-fixation, the larvae were permeabilized with 0.5% Triton in 1X PBS and washed in deionized water for 1 hour. Blocking was performed with 2% BSA in 0.5% Triton/PBS. Anti-GFP (chicken-A10262), anti-mCherry (mouse-A10262), and Vcam1b (Cat. No. DZ01199-1; Boster Bio) were used in 1:200 dilution overnight at 4°C. Anti-Chicken-488 (A11039), Anti-mouse-568 (A10037) and Anti-rabbit-647 (ab150079) were used as a secondary antibody at 1:1000 dilution overnight at 4°C. Imaging was performed using a Yokogawa spinning disk and Nikon inverted Ti microscope.

### Whole mount *in situ* hybridization (WISH)

Embryos were stage-matched and fixed overnight at 4°C in 4% paraformaldehyde (PFA). WISH was performed as previously described (Thisse and Thisse, 2008b) with the following exceptions: before permeabilization with proteinase K, embryos were bleached to remove pigmentation, and 0.2% glutaraldehyde was added to 4% formaldehyde at the post-fixation step. Antisense RNA probes were *cmyb* (Liao et al., 1998) and *vcam1b*. The *vcam1b* probe for was generated using a plasmid containing *vcam1b* cDNA (ZGC vcam1) purchased from Horizon (clone ID: 8149942). The plasmid was digested using EcoRI-HF, NotI-HF, and FspI (NEB, R3101, R3189, R0135). The sample was analyzed on an agarose gel, and the band size of ∼3,000 bp was extracted using the QIAquick Gel Extraction Kit (Cat no. 28704). The pBSKSII plasmid was digested with EcoRI-HF and NotI-HF. The digested plasmid was purified with the QIAquick PCR Purification Kit (Cat no. 28104) and treated with alkaline phosphatase. The PCR fragment and pBSKSII plasmid were ligated utilizing T4 ligase in a 2:1 ratio (Promega, cat no. M1801) and transformed into TOP10 competent cells (Invitrogen, C404003). The ligation was confirmed by digestion. For probe synthesis, the pBSKSII-vcam1 plasmid was digested with EcoRV (NEB, R0195), followed by inverse *in vitro* transcription with T7 transcription enzyme. The synthesized product was then purified and stored at −80°C for further use.

### Intravascular antibody injections in the embryo

Embryos were screened at 50 hpf for fluorescent markers (*mpeg1:GFP* and *kdrl:mcherry*) and injected by retro-orbital injections with a 1 nl containing either Vcam1b-647 antibody (final concentration 0.4 mg/mL) or an isotype control IgG-647 (ab150079, Abcam; 0.4 mg/mL). Solutions were co-injected with Dextran, Cascade Blue, 10,000 MW, Anionic, Lysine Fixable dye (Invitrogen; Cat. No. D1976; 5 mg/mL) to verify successful delivery into the circulatory system. 2 hours post-injections, larvae were mounted in 1% LMP agarose in E3 media, kept in E3 media with tricaine for live imaging. Live imaging was performed using a Yokogawa spinning disk and Nikon inverted Ti microscope

### Flow cytometry and sorting

Embryos from the indicated transgenic lines were euthanized with tricaine at 54 hpf. Single-cell suspensions were prepared by adapting the protocol previously described (Bresciani et al., 2018). Briefly, embryos were dissociated in PBS containing 2% FBS using a combination of mechanical pipetting, and enzymatic digestion at 35°C. The cell suspension was filtered through a 70 µm cell strainer, followed by centrifugation at 300x g. Cells were resuspended in 1xPBS supplemented with 2% FBS. For immunostaining, cells were incubated with anti-Vcam1b-iFluor-647 antibody (1:50; Cat. No. DZ01199-1-iFluor647; Boster Bio) or Goat Anti-Rabbit IgG-Alexa Fluor 647 isotype control (1:50; Cat. No. ab150079; Abcam) for 30 minutes at RT in the dark. Cells were counted with a hemocytometer and trypan blue. Flow cytometry analysis was performed using a BD LSRII Fortessa flow cytometer. Cell sorting was performed with a BD FACSAria II and dead cells were excluded using DAPI (4′,6-diamidino-2-phenylindole).

### Single cell RNA sequencing

Approximately 100 *casper* and 100 *cxcl12:dsRed*+ embryos were dissociated, as described above, for cell sorting of anti-Vcam1b-iFluor-647+ cells. Sorted cells from both backgrounds were pooled and then split into 2 technical replicates for loading onto a Chromium Single Cell Controller (10x Genomics) to generate single-cell gel beads in emulsion (GEMs) by using Single Cell 3’ Library and Gel Bead Kit V3.1 (10x Genomics, 1000121). Cell lysis and RNAs barcoding were accomplished through reverse transcription in individual GEMs. Single-cell RNA-seq libraries were prepared using Single Cell 3’ v3.1 Reagents Module 2 (10x Genomics, 1000121) following the manufacturer’s instructions. A MiSeq balancing run was performed, followed by paired end sequencing on an Illumina NovaSeq X Plus.

### Single cell RNA-seq analysis

CellRanger (5.0.0) was used to generate gene expression containing unique molecular identifies (UMIs) matrix using the default parameters. We used a custom zebrafish reference genome (custom_GRCz11_reporters) that included cDNA sequences for expressed transgenes (e.g., EGFP and mCherry). Samples were then further filtered and clustered in Seurat (Stuart et al., 2019). We flagged and removed cells with low detected features or high percent of mitochondrial reads. The threshold we used was 2 median absolute deviations (MADs) from the median for mitochondrial reads and the cutoff was 1.9%. For detected features we used a threshold of 1 MAD as a more stringent cutoff. DoubletFinder was used to remove a small number of high-confidence doublets (McGinnis et al., 2019). At 1 MAD the cell counts in technical replicates were 1,331 (vcam1b_1) and 1,371 (vcam1b_2). Technical replicates were highly similar (median UMI count within 0.5%) so no additional normalization was performed. Multiple clustering algorithms were tested (Partitions, Leiden, Louvain) and Leiden clusters were presented in Figure 3. Cluster annotation was manually curated after using the Seurat label transfer algorithm (Stuart et al., 2019) and publicly available data (Farnsworth et al., 2020).

### Reanalysis of published sorted cell population bulk RNA-seq datasets

When provided by study authors, we used gene expression values normalized for library size and gene-of-interest length. For studies in which only unnormalized read counts were available, read counts were normalized for library depth with DESeq2 (Love et al., 2014) and for estimated gene length. Where only DESeq2 normalized read counts were made available, expression values were additionally normalized for estimated gene length. Gene lengths in kilobases were estimated by calculating the number of base pairs covered by at least one annotated exon for that gene (present in GRCz11 dataset from Ensembl release 112) and dividing by 1000. Reanalyzed sorted EC bulk RNA-seq datasets: vein-specific at 38 hpf (GSE152759: *Nr2f2+/Tg(fli1a:egfp)+*) (Lawson et al., 2020); blood and vasculature at 38 hpf (GSE152759: *Tg(fli1a:egfp*)+) (Lawson et al., 2020); vasculature at 120 hpf (GSE119718: *Tg(kdrl:HRAS-mCherry)+* and negative) (Whitesell et al., 2019). Reanalyzed sorted mesenchymal cell bulk RNA-seq datasets: smooth muscle cells at 96 hpf (GSE119718: *Tg(foxc1b:EGFP)+;Tg(acta2:mCherry)+* and negative) (Whitesell et al., 2019); vascular mural cells at 96 hpf (GSE119718: *Tg(acta2:EGFP)+* and negative); epidermis, cartilage, pericytes, and vascular smooth muscle cells cells at 120 hpf (GSE152759: *TgBAC(pdgfrb:citrine)+* and negative) (Lawson et al., 2020). Reanalyzed sorted macrophage bulk RNA-seq datasets (renormalized together): 28 and 50 hpf (GSE149789: *Tg(mpeg1:GFP)+*) (Kuil et al., 2020); 72 hpf (GSE93818: *Tg(mpeg1:mCherry)+* and negative) (Theodore et al., 2017); 120 hpf (GSE78954: (*Tg(mpeg1:mCherry)+*) (Rougeot et al., 2019); 144 hpf (GSE78954: *Tg(mpeg1:GAL4;UAS:Kaede)+* and (*Tg(mpeg1:mCherry)+*) (Rougeot et al., 2019). Additional details in Extended Data Table 1.

### Whole mount fluorescent in situ hybridization (FISH) and immunofluorescence (IF)

Zebrafish embryos from *mpeg:GFP*, *kdrl:mCherry*, and *ET37:GFP* transgenic lines were fixed overnight at 4°C in 4% formaldehyde in 1x PBS at 54 hpf. Following fixation, embryos were rinsed with PBST (0.1% Tween-20 in PBS), then dehydrated and stored in 100% methanol at −20°C.

Embryos were rehydrated and RNAscope™ Multiplex Fluorescent Reagent Kit v2 (Advanced Cell Diagnostics, Cat. No. 323100) was used according to the manufacturer’s instructions with several modifications. Embryos were incubated in 3% H_2_O_2_ for 10 min at RT, followed by washes with PBST, and then treated with RNAscope Protease III (Cat. No. 322337) for 5 min at RT and subsequently washed three times with 0.01% PBST (0.01% Tween-20 in PBS). A prehybridization step was included by incubating embryos at 40°C for at least 2 h in RNAscope probe diluent (Cat. No. 300041). The vcam1b-C3 probe (Cat. No. 445681-C3) was then applied, and embryos were incubated overnight at 40°C. All subsequent wash steps were performed with 0.01% Tween-20 in 0.2X SSCT, three times for 15 min each. Post-probe incubation, a postfix treatment with 4% PFA was conducted for 20 min at RT. All 40°C incubation steps were performed in a water bath. Following the RNAscope protocol, embryos were washed three times with PBST. Blocking was performed for 1 h (2% BSA in PBST). Embryos were incubated overnight at 4°C with anti-GFP antibody (1:500; chicken, Invitrogen, Cat. No. A10262) or anti-mCherry antibody (1:500; Mouse, Abcam, Cat No. ab125096). Embryos were washed before secondary antibody incubation. Secondary antibodies anti-chicken-Alexa 647 (1:1000; Goat, Abcam, Cat No. ab150175) or anti-mouse-Alexa-647 (1:1000; Donkey, Invitrogen, Cat No. A31571) were incubated overnight at 4°C. Embryos were washed with PBST and nuclear staining was performed using DAPI. Before imaging, embryos were washed with 0.01% Tween-20 in PBS and mounted in 1% low melting point agarose (Invitrogen; Cat. No. 16520) in PBS for confocal imaging. Imaging was performed with a Nikon A1R inverted Ti2 confocal microscope and a Nikon 20x/Plan Apo/Air/NA=0.80/WD=1mm objective lens.

### Image analysis

Images acquired from RNAscope and immunohistochemistry were analyzed using Imaris software. *Vcam1b* expression from RNAscope data was converted into 3D spots using the Imaris Spot Detection tool. The threshold for positive signal detection was determined for each image based on the center pixel intensity value. Cell surfaces were subsequently defined based on the immunohistochemical staining using the Imaris Surface tool. The number of *vcam1b* spots located within 0 µm of the defined cell surfaces were quantified for our analysis. Whole mount *in situ* hybridization images were quantified using Fiji/ImageJ and an unbiased scoring approach (Dobrzycki et al., 2020).

### Software

Statistical analyses and plots were done using R (Version 4.4.0) or GraphPad Prism (Version 10.5.0). Image analysis was performed with Imaris (Version 10.2.0; Oxford Instruments), Nikon NIS-Elements AR (Version 5.20.02), and Fiji/ImageJ. Flow cytometry data was analyzed with FlowJo (Version 10.9).

## Online supplemental material

This manuscript contains 3 supplementary figures, 1 supplementary table, and 4 supplementary movies. Figure S1 “Confirmation of *vcam1b* expression and function in CHT hematopoiesis” contains reanalysis of published data, additional description of *vcam1b* gene expression, mutant, and polyclonal anti-Vcam1b antibody. Figure S2 “Intravascular injection of conjugated antibodies produces broad non-specific staining in the CHT” highlights challenges of this technique with live imaging of injected embryos. Figure S3 “UMAP plots of published scRNA-seq data showing *vcam1b* is expressed in ECs and MSCs of embryonic and larval zebrafish”. Table S1 is related to Figure 4 and includes normalized counts of selected genes after reanalysis of published sorted bulk RNA-seq datasets from specific cell populations. Movie 1 relates to Figure 1 and shows *Runx:GFP*+ HSPCs in the CHT of a WT or Het (*itga4^cas005^*^/+^) *kdrl:mCherry*+ embryo. Movie 2 relates to Figure 1 and shows *Runx:GFP*+ HSPCs in the CHT of a homozygous mutant (*itga4^cas005/cas005^*) *kdrl:mCherry*+ embryo. Movie 3 relates to Figure S2 and shows time-lapse live imaging of the CHT of a *mpeg1:GFP;kdrl:mCherry* embryo after intravascular injection with Vcam1b-647 antibody. Movie 4 relates to Figure S2 and shows time-lapse live imaging of the CHT of a *mpeg1:GFP;kdrl:mCherry* embryo after intravascular injection with IgG-647 antibody.

## Data availability

scRNA-seq data is available in NCBI GEO (Series: GSE304537). Additional information, data, and requests for resources and reagents should be directed to the corresponding author, O.J.T.

## Acknowledgments

We thank: Drs. Samuel J. Wattrus and Leonard I. Zon for critical reading of the manuscript; Carbone Cancer Center Flow Cytometry Laboratory (supported by P30 CA014520, BD LSR Fortessa (1S10OD018202-01), BD FACS AriaII (1S10RR025483-01)); Harrison Pantera and Sandra Splinter BonDurant at UW-Madison Biotechnology Gene Expression Center (RRID:SCR_017757); UW-Madison Next Gen DNA Sequencing Core; UW-Madison Bioinformatics Resource Center (RRID:SCR_017799); Lance Rodenkirch at UW-Madison Optical Imaging Core facility. E.J.H. was supported by a National Institutes of Health (NIH) National Institute of Diabetes and Digestive and Kidney Diseases (NIDDK) grant (K01DK111790), a Career Investment Award from the BU Department of Medicine and an AABB Early Career Scientific Research Grant. B.W.B. was supported by a NIH NIDDK grant (R01DK128238). O.J.T was supported by the NIH National Heart, Lung, and Blood Institute (NHLBI) grants (R01HL174965, R01HL142998, R56HL142998), an American Society of Hematology (ASH) Bridge Grant Award, and the Department of Cell and Regenerative Biology at UW-Madison.

---

**Figure S1.**
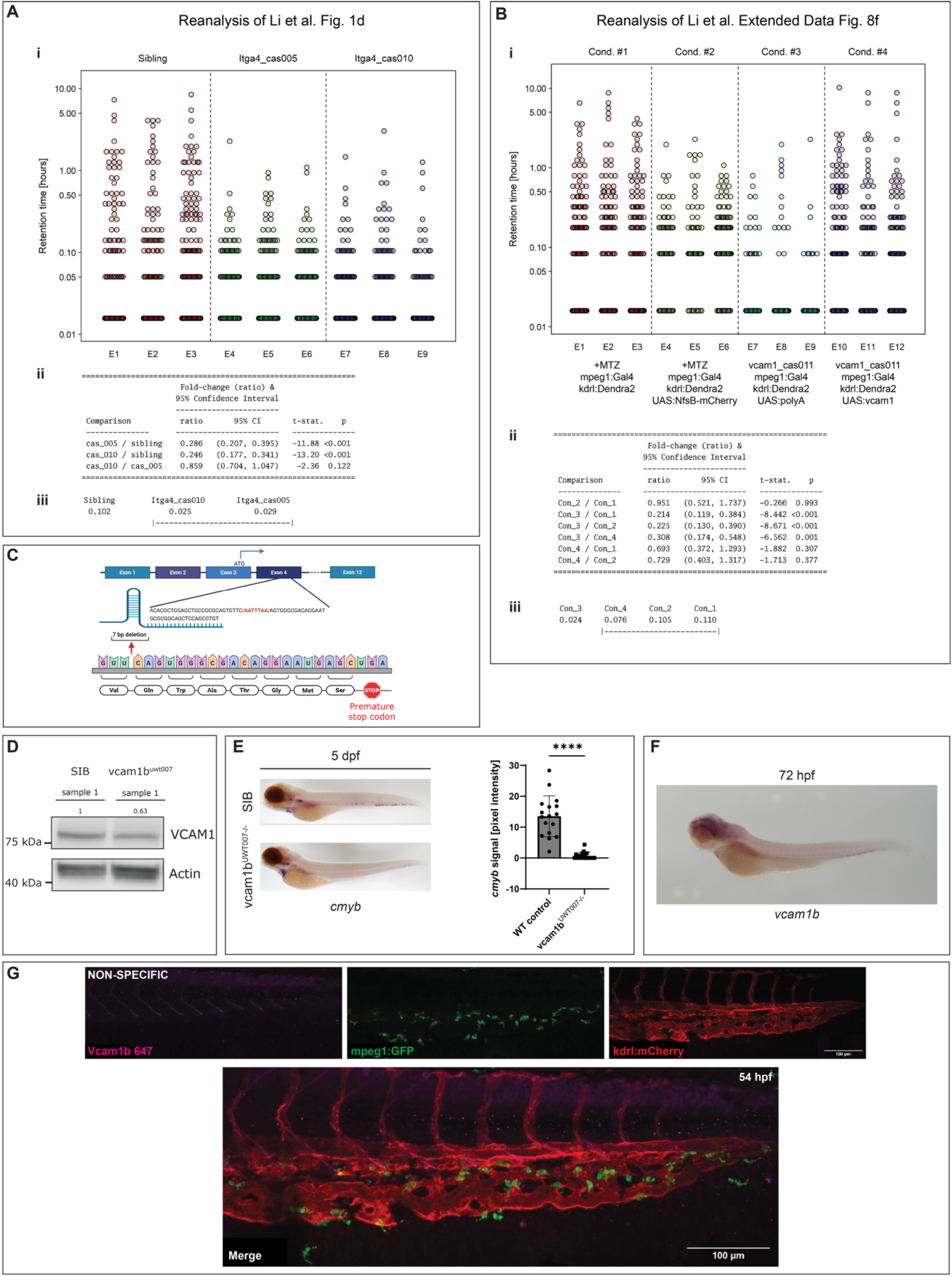
Confirmation of *vcam1b* expression and function in CHT hematopoiesis. **(A)** Reanalysis of HSPC retention times in *itga4* mutants (previously published raw data; Figure 1d (Li et al., 2018)). **(i)** This figure involved 3 genotypes (Sibling, *itga4^cas005^*, and *itga4^cas010^*) with n=3 biological replicates per genotype. Retention time ranged from 0.015–9.05 hours with 58.8% of all readings (743/1263) fixed at the minimum. The 75^th^ percentile of retention time (for all readings) was 6 minutes and the observed fraction of readings in each treatment with retention times <6 minutes (0.10 hrs) was 47.4%, 80.9%, and 85.8% for sibling, *itga4^cas005^*, and *itga4^cas010^*, respectively. Plot shows the distribution of readings within each biological replicate within a given genotype. Vertical axis has log scaling. A linear mixed-effect model (performed on log-transformed retention times with embryo as the random effect and treatment as the fixed factor) revealed significant differences (p<0.001) in retention time for each *itga4* mutant allele relative to sibling. The geometric mean (GM) retention time for the *itga4* mutant alleles together is roughly 73% less than the GM retention time for the sibling condition (i.e., ∼71% lower for *itga4^cas005^* and ∼75% lower for *itga4^cas010^*). There was no evidence (p=0.122) to suggest any difference between the two *itga4* mutant alleles (*itga4^cas005^* vs *itga4^cas010^*). **(ii)** All t-statistics are based on 6 degrees of freedom (df) and p-values adjusted using Tukey’s method. **(iii)** Ordering of experimental conditions according to GM retention time. Conditions sharing a common underline do not differ at the 0.05 level (Tukey’s method) and form a homogeneous group. Conclusions are similar if one considers the proportion of retention times that are <6 minutes. Both *itga4* mutant alleles strongly differ from the sibling condition (p<0.001 for each), while there is no appreciable difference between the two alleles of *itga4* (p=0.351). **(B)** Reanalysis of HSPC retention times in a *vcam1b* mutant (Cond. #3; previously published raw data; Figure 8f (Li et al., 2018)). **(i)** n=3 embryos for each experimental condition (#1–4), with ∼95 recorded cells/embryo. Retention times ranged from 55 seconds to 9.5 hours. The minimum retention time of 55 seconds was observed in: 37% of cells from Cond. #1; 20% of cells from Cond. #2; 84% of cells from Cond. #3; 44% of cells from Cond. #4. GM retention times for Cond. #1–4 were 0.11, 0.10, 0.02, and 0.08 hours, respectively. Cond. #3 is just barely above the observed 55 second minimum (due to only 16% of retention times ever being longer than 55 seconds), while Cond. #1 and #2 are similar. Of the six possible pairwise comparisons, only the three involving Cond. #3 are significant. There is no difference (t_8_ = −0.266, p = 0.993) between Cond. #1 and #2 with respect to GM retention time, with times being ∼5% lower for Cond. #2 relative to Cond. #1 (i.e., ratio ≈ 0.951). GM retention time for Cond. #3 is ∼80% lower than the corresponding GM retention time for each of Cond. #1 and #2 and is ∼70% lower than the GM retention time for Cond. #4. The table below shows all pairwise comparisons with fold changes (ratios) and their supporting 95% confidence intervals (CIs). **(ii)** All t-statistics are based on 8 degrees of freedom (df) and p-values adjusted using Tukey’s method. **(iii)** Ordering of experimental conditions according to GM retention time. Conditions sharing a common underline do not differ at the 0.05 level (Tukey’s method) and form a homogeneous group. **(C)** Schematic illustrating the CRISPR-Cas9 strategy used to generate the *vcam1b^uwt007^* mutant we reported previously (Bornhorst et al., 2024). The edit resulted in a 7-bp deletion and a premature stop codon. **(D)** Representative western blot of sibling (SIB) and *vcam1b^uwt007^* mutant brain samples. Upper panel shows Vcam1b protein expression, lower panel shows actin loading control. **(E)** Left: Representative WISH images of HSPC marker *cmyb* at 5 dpf that was reduced in *vcam1b^uwt007^*mutants compared to SIB controls. Right: Quantification of the number of *cmyb* expressing cells in the CHT at 5 dpf. n=17 SIB controls vs n=19 *vcam1b^uwt007^*mutants; Unpaired t test; ****p<0.0001. Anterior at left, dorsal at top. **(F)** Representative WISH expression pattern of *vcam1b* at 72 hpf. Expression in brain, heart, and CHT. Note: to orient anterior at the left, the image was transformed by reflection on the vertical axis. **(G)** Whole mount IF of fixed 54 hpf embryo with polyclonal anti-Vcam1b antibody (magenta; upper left) is NON-SPECIFIC and NOT REPRESENTATIVE of endogenous mRNA expression pattern shown in (F). Representative IF for *mpeg1:GFP* (anti-GFP; green; upper middle) and *kdrl:mCherry* (anti-mCherry; red; upper right). Lower enlarged image shows 3 channels merged. Scale bars: 100 microns. Note: to orient anterior at the left, the image was transformed by reflection on the vertical axis.

**Figure S2.**
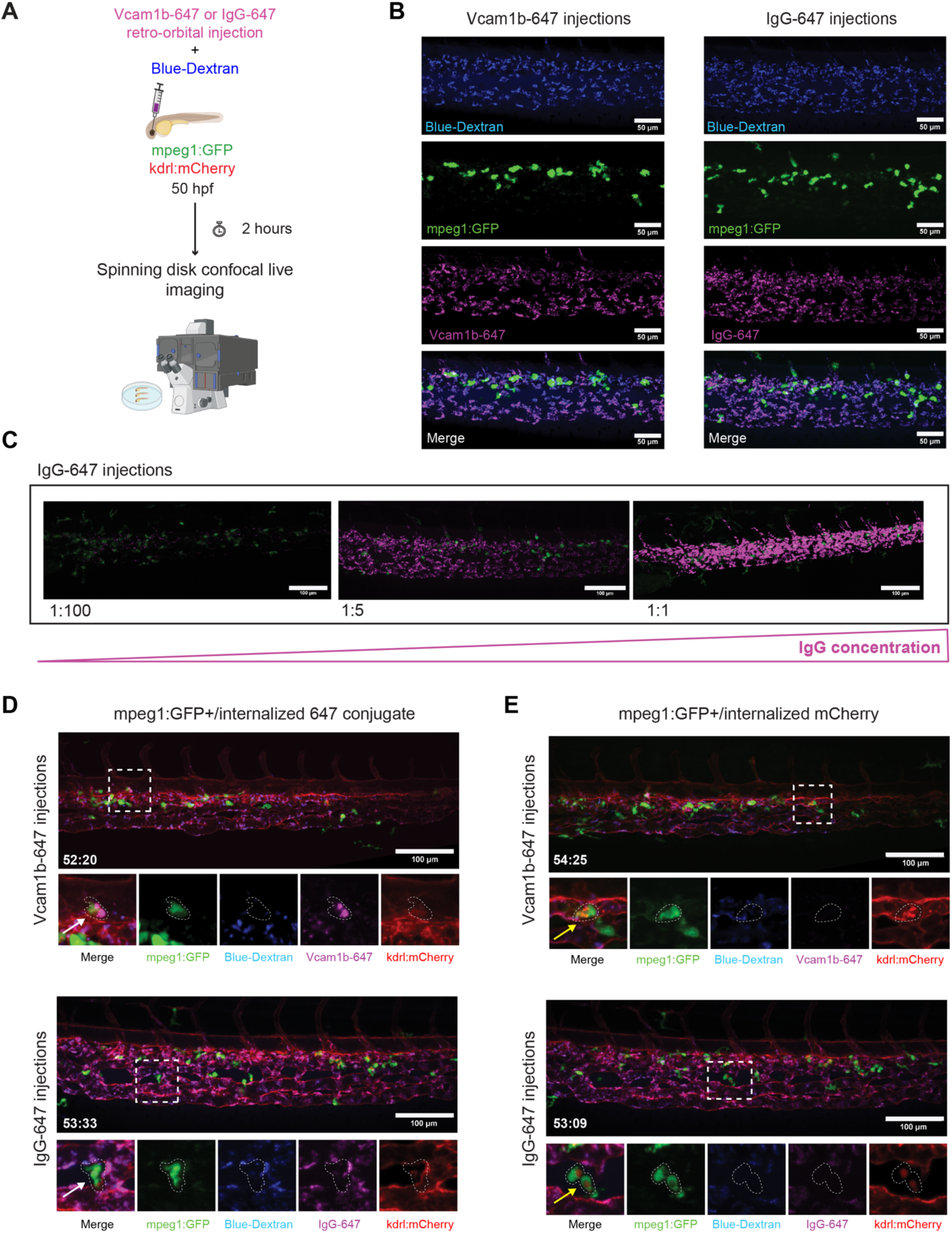
Intravascular injection of conjugated antibodies produces broad non-specific staining in the CHT. **(A)** Diagram showing experimental design. Vcam1b-647 conjugated antibody or IgG-647 isotope control, together with blue dextran, were injected retro-orbitally into *mpeg1:GFP*+ transgenic embryos at 50 hpf. Live imaging of the CHT was performed 2 hours after injection. **(B)** Representative images of the CHT in live embryos. 3 channels: blue dextran (blue); mpeg1:GFP (green); 647 conjugate (magenta). Bottom is merge. Left: Vcam1b-647 injected. Right: IgG-647 injected. Vcam1b-647 conjugated antibody or IgG-647 isotope control have similar NON-SPECIFIC labeling in the CHT. Scale bar is 50 microns. **(C)** Concentration-dependent NON-SPECIFIC signal after injection of IgG-647 isotype control (1:100, 1:5, 1:1 dilutions) into *mpeg1:GFP*+ embryos. **(D-E)** Representative single frames from time-lapse movies of *kdrl:mCherry;mpeg1:GFP* embryos injected with Vcam1b-647 conjugated antibody or IgG-647 isotope control and blue dextran. *mpeg1:GFP+* macrophages carrying internalized **(D)** 647 conjugates or **(E)** debris from remnants of *kdrl:mCherry*+ ECs injected with Vcam1b-647 (upper) or IgG-647 (lower).

**Figure S3.**
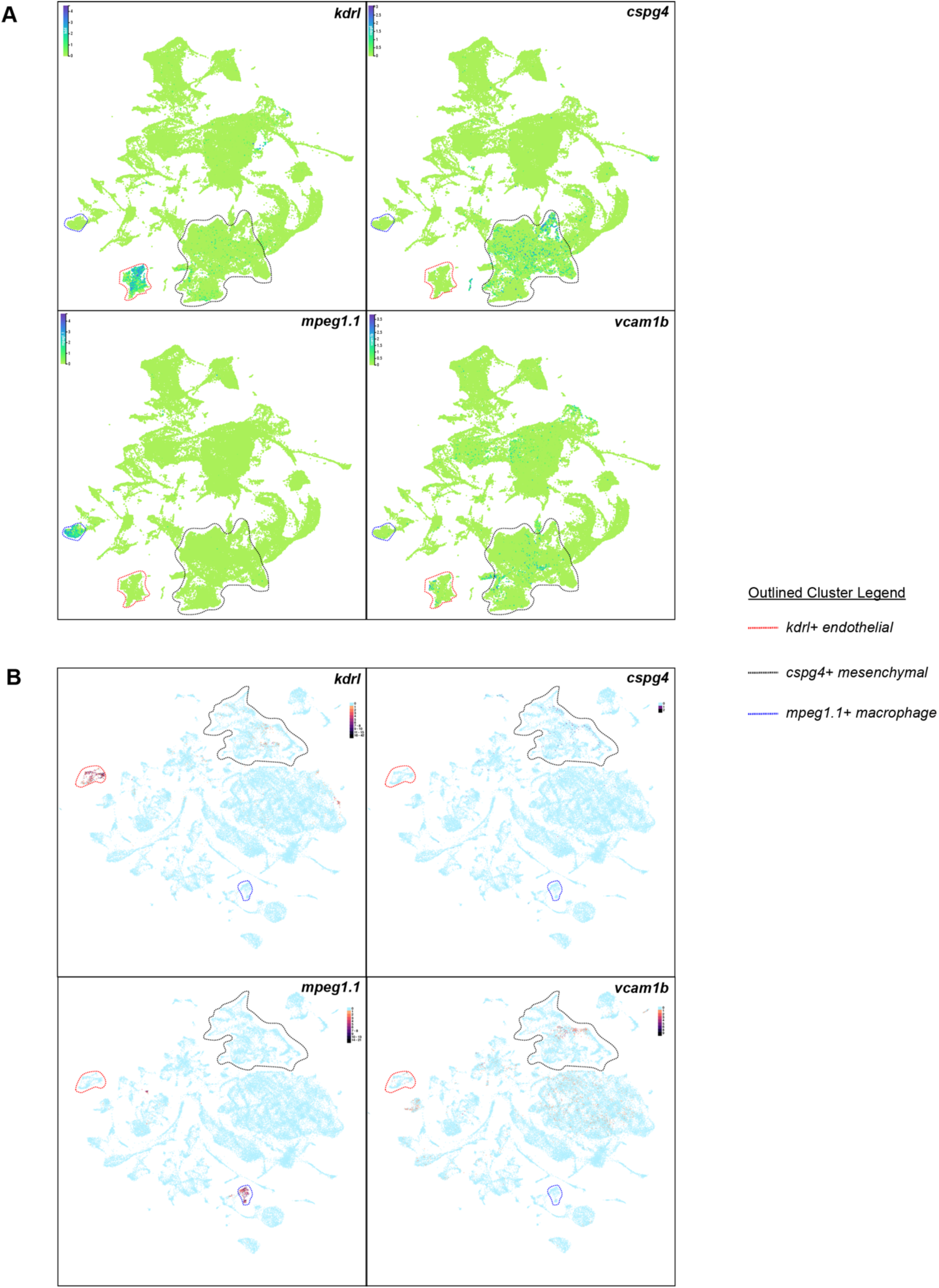
UMAP plots of published scRNA-seq data showing *vcam1b* is expressed in ECs and MSCs of embryonic and larval zebrafish. **A)** Full Zebrahub (Lange et al., 2023) dataset with stages from 10 hpf to 10 dpf. EC-specific *kdrl* and MSC-expressed *cspg4* mark clusters outlined in red and black, respectively. These clusters also have cells that express *vcam1b*. In a cluster marked by macrophage-specific *mpeg1.1* (outline in blue) there is no expression of *vcam1b*. **B)** In another large publicly available zebrafish scRNA-seq dataset with stages 1, 2, and 5 dpf (A Single-Cell Transcriptome Atlas for Zebrafish Development displayed with the UCSC Cell Browser (Farnsworth et al., 2020; Speir et al., 2021)), we observed similar results as in (A).

